# Sorting single T cells based on secreted cytokines and surface markers using hydrogel nanovials

**DOI:** 10.1101/2022.04.28.489940

**Authors:** Doyeon Koo, Robert Dimatteo, Sohyung Lee, Joseph de Rutte, Dino Di Carlo

## Abstract

Immune cell function is intrinsically linked to secreted factors which enable cells to communicate with neighboring or distant cells to coordinate a response. The ability to secrete cytokines also can help define the population of cells with therapeutic potential in emerging cell therapies, such as chimeric antigen receptor (CAR)-T cell therapies. Polyfunctional cells that can secrete more than one cytokine have been found to play an outsized role in therapeutic efficacy. While there are a variety of techniques to analyze cellular secretions from individual polyfunctional cells, there are no widely-available approaches to sort viable cells based on this phenotype. Here, we apply lab on a particle technology to the analysis and sorting of T cells based on a combination of secreted factors, interferon gamma (IFN-γ), tumor necrosis factor alpha (TNF-α) and interleukin 2 (IL-2) and surface markers (CD8+ and CD4+). Cells are selectively loaded into the antibody-functionalized cavity of micro-hydrogel particles, called nanovials, where secreted cytokines are captured and fluorescently stained. By leveraging standard fluorescence activated cell sorters and using fluorescence pulse area/height information we can distinguish between fluorescence signals on the nanovial cavities and on cells, and are able to process greater than 1 million nanovials in one hour of sorting. The frequency of multi-cytokine secreting cells was correlated with surface marker expression, and biased towards CD4+ T cells. CD8+ cells that secreted more than one cytokine, were biased towards IFN-γ and TNF-α with fewer CD8+ cells secreting IL-2. The majority of cells with a polyfunctional phenotype that were sorted remained viable and regrew following sorting. This nanovial cytokine secretion assay can be applied to sort antigen-specific T cells or CAR-T cells based on their functional engagement with cognate antigens or peptide-major histocompatibility complexs (MHCs), enabling discovery of functional CARs or T cell receptors and deeper investigation into the molecular underpinnings of single T cell function.

## Introduction

A key function of immune cells is the ability to secrete communication factors that help coordinate a response to a pathogen. Communication factors, such as cytokines, extend beyond the local contact-based influence of surface expressed proteins to signal to many neighboring cells, leading to an outsized importance for the function of cells. Other enzymes and pore-forming secreted factors are critical to other effector functions of immune cells, such as the ability to kill target cells that may be infected or malignant. Measurement of the secretions from single immune cells and sorting based on these secretions provides biological insight into the ultimate function of immune cells and can help researchers engineer immune cells for treating disease.

In the future, engineered cell therapies will be a pillar of medicine, along with molecular (drugs and proteins) and genetic (gene therapy) interventions. In the last few years, there has been particular success in the use of engineered immune cell-based therapies in treating hematologic malignancies, including recent FDA approvals of chimeric antigen receptor (CAR)-T-cell therapies. Unfortunately, complete remission rates can vary from 20-90% and treatments, as of yet, have failed to show efficacy for solid tumors such as breast, colon, and pancreatic cancers^1–3^. The biological basis behind this variation in outcomes is still not well understood and provides a significant barrier in our ability to standardize across treatment approaches and define quality control metrics between batches of the same treatment^4^. However, a growing body of literature has alluded to certain T cell functional properties such as high proliferative potential and propensity to secrete multiple cytokines simultaneously as key drivers of response^5–7^. Therefore, a more thorough understanding of the functional heterogeneity in T cell populations may assist in the next generation of cancer immunotherapies. In fact, recent single-cell screens have highlighted an astonishingly high level of functional diversity from T cells isolated from the same patient and bearing the same panel of surface markers, with only a small highly active subset of cells driving responses to immunological challenge^8–10^. New tools to sort based on cell function, such as level and type of secreted cytokine, can enable identifying gene expression signatures associated with functional responses, uncovering cell surface markers that are more descriptive of optimal functional phenotypes, and discovery of T cell receptors/CARs that move beyond high affinity to elicit robust functional responses. Ultimately, these tools can be used to directly sort optimal therapeutic cells based on function, leading to more effective cellular medicines.

Notably, widespread tools to analyze and select populations of therapeutic cells based on functional properties, such as secreted products, are lacking. The most basic approach, intracellular cytokine staining (ICS), can have high throughput by leveraging flow cytometry, but requires fixation and permeabilization, rendering cells non-viable for subsequent culture or functional assays ^11^. Techniques like ELISpot can analyze the secretions from viable cells, however, like ICS, this is a terminal assay and cells cannot be sorted following the measurement to select high or low secretors. Cytokine capture probes, commercialized by Miltenyi Biotec, translate secreted proteins into fluorescent signals bound to cell surfaces, but they are susceptible to crosstalk and loss of dynamic range due to secreted cytokines diffusing and adhering to neighboring non-active cells^12–14^. Further, cytokines which may have surface and secreted forms, like tumor necrosis factor alpha (TNF-α) cannot be separately measured. Microfluidic analysis systems lend themselves well to functional screening applications because of their ability to precisely manipulate small liquid volumes to isolate cells into microwells or water-in-oil emulsions to concentrate secreted proteins and prevent crosstalk. Companies such as Isoplexis and Berkeley Lights have commercialized several all-in-one microfluidic platforms requiring minimal user input, but the specialized nature of such tools, combined with requirements for capital equipment purchases ($100,000 - $2,000,000 base cost), hinder adoption and access by research groups and small companies^8,15^. In addition, the approaches are limited to analyzing thousands to tens of thousands of cells, much less than standard flow cytometry that can analyze thousands of events per second. Although the Isoplexis approach uses microfluidic barcoding which enables analyzing tens of cytokines, the cells cannot be recovered after the assay, similar to ICS or ELISpot.

Recently, we reported a microfluidics-free approach to confine cells into sub-nanoliter compartments using hydrogel particles with bowl-shaped cavities, which we call nanovials, and capture secreted molecules on the nanovial surfaces. We applied the nanovial technology to isolate and sort B cells based on their production of antibodies or the production of antibodies with specific affinity for an antigen^16^. Our previous work also showed that by linking gelatin only to the inner cavity of a nanovial we could localize the capture of cells and their secretions in the cavity and further reduce secreted antibodies from spreading to neighboring nanovials^17^.

Here, we have applied the localized gelatin nanovial technology for high-throughput analysis and sorting of individual primary T cells based on secreted cytokines using a fluorescence activated cell sorter (FACS) (Figure 1). Nanovials are functionalized with cytokine capture antibodies (anti-IFN-γ, anti-TNF-α, anti-IL-2) and cell binding motifs (anti-CD45). T cells are loaded into the cavities of nanovials and, following activation, secrete cytokines which are also captured in the cavity of the nanovial with minimal crosstalk. Captured cytokines are labeled with a fluorescent detection antibody while cells are labeled with viability dyes or additional surface markers. Cytokine binding to nanovials could be easily distinguished from non-specific cell binding through use of fluorescence peak area and height metrics associated with staining of the larger nanovials. We demonstrate the ability to analyze and sort viable cells based on four markers simultaneously; two surface markers (CD4 and CD8) and two secreted proteins (IFN-γ and TNF-α or IL-2). Cells are sorted based on the secretion levels using a commercial FACS and remain viable and can be re-cultured after sorting. Up to 1 million nanovials could be analyzed and sorted within 1 hour.

**Figure 1.**
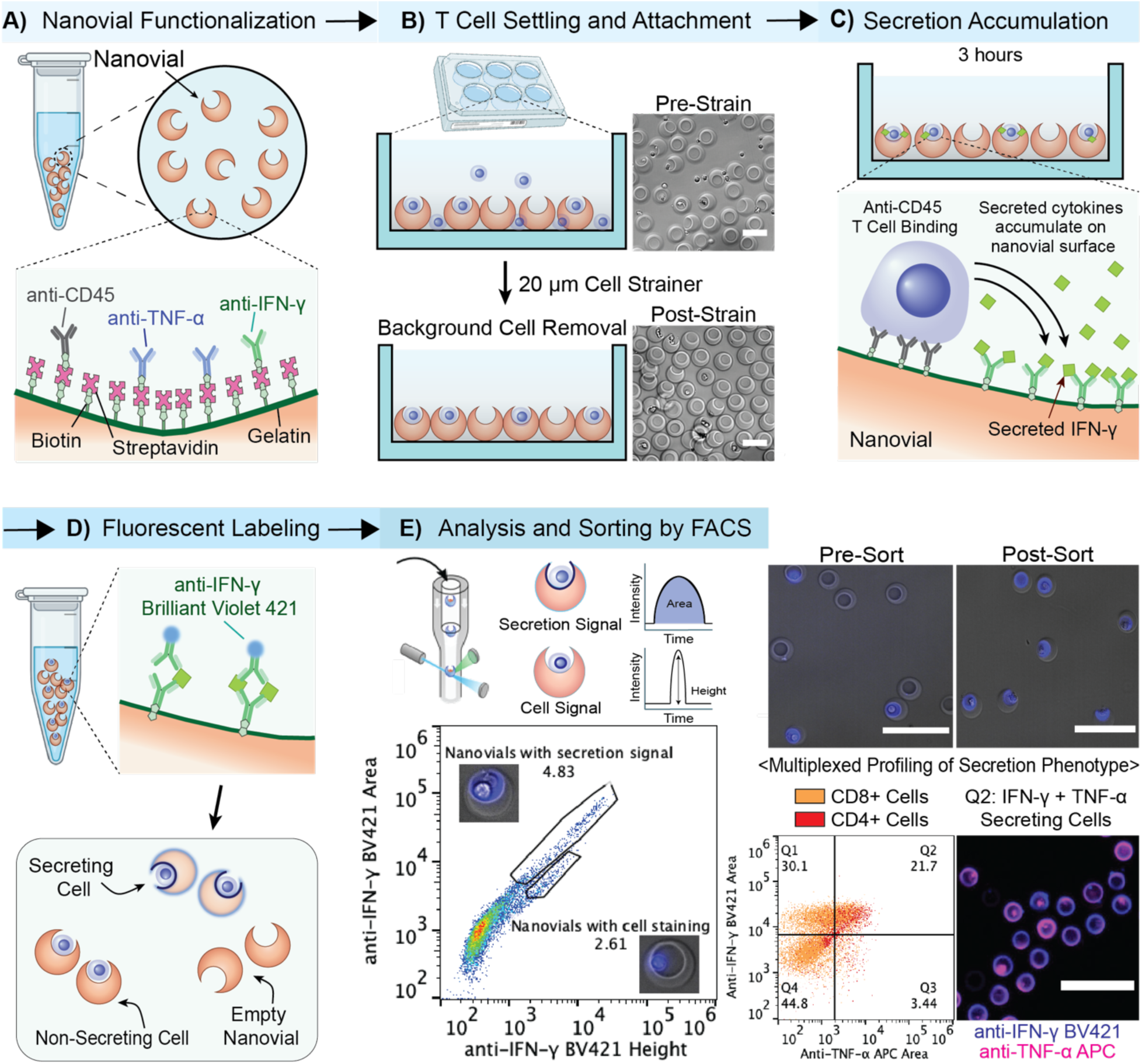
Overview of high-throughput analysis and sorting of individual T cells based on secreted cytokines using nanovial technology. A) Nanovials are functionalized with biotinylated cytokine capture antibodies and cell binding motifs via streptavidin-biotin chemistry. B) T cells are loaded into the cavities of nanovials in a well plate and any unbound cells are removed using a cell strainer. Scale bars represent 50 μm. C) T cells are activated for 3 hours and secreted cytokines are captured in the cavity of the nanovial. D) Captured cytokines are labeled with a fluorescent detection antibodies and cells may be labeled with additional surface markers. E) Cytokine binding to nanovials can be distinguished from non-specific cell staining using a combination of fluorescence peak area and height signals associated with staining of the larger nanovials. Cells are sorted based on the secretion signal as well as cell surface markers using a commercial FACS. Scale bars represent 100 μm.

## Results

### Fabrication and functionalization of nanovials

Using an aqueous two-phase system combined with droplet microfluidics and hydrogel chemistry, we fabricated cavity-containing hydrogel microparticles. With some modifications on our lab’s previously reported study^16,17^, we used a flow-focusing device to generate millions of monodisperse polyethylene glycol (PEG)-based nanovials with the inner cavity selectively coated with biotinylated gelatin (Figure S1.A). To accommodate human T cells with diameters of ∼10 μm we fabricated uniform nanovials with an average diameter of 35 μm (CV = 5.1%) and average cavity of 21.2 μm (CV = 7.2%). Nanovials are functionalized with biotin, reacted with streptavidin and then linked to a number of different biotinylated antibodies to selectively isolate cells based on surface expression and capture secreted molecules. For T cell secretion assays we decorate nanovials with anti-IFN-γ and anti-TNF-α or anti-IL-2 antibodies along with anti-CD45 antibodies to selectively adhere T cells into the cavity of nanovials. In total, 2 or 3 antibodies were linked to the nanovials through streptavidin-biotin noncovalent interactions. We found that the optimum ratio between anti-CD45 capture antibodies and cytokine secretion capture antibodies was 1:1 (140 nM each) for single cytokine analysis and 1:1:1 (140 nM each) for multiplexed probing, which allowed for T cell loading as well as signal from recombinant cytokines down to 10 ng/mL (Figure S1.B, Figure 2C).

**Figure 2.**
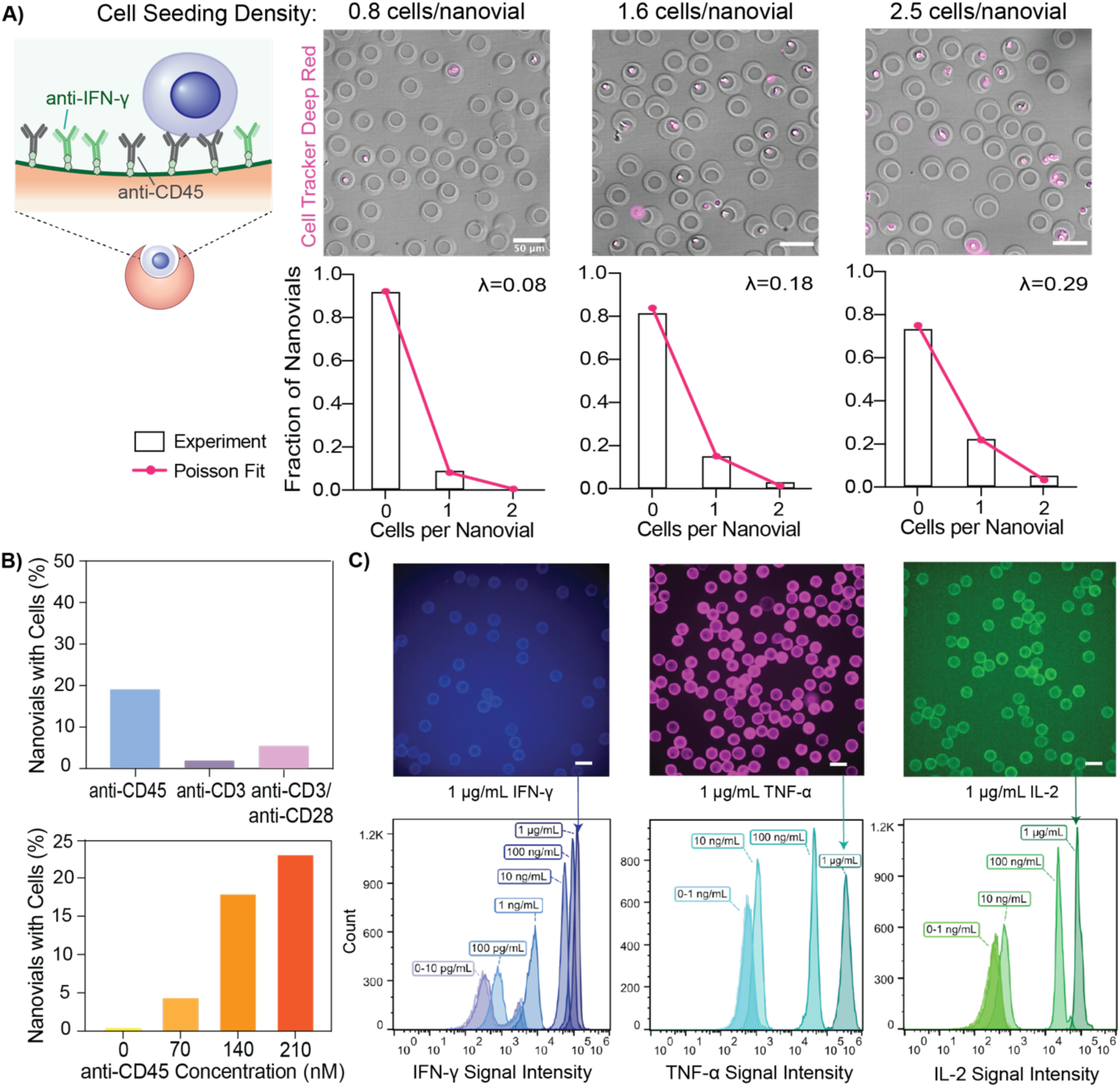
Loading statistics and dynamic range of nanovials. A) T cells were loaded onto anti-CD45 and anti-IFN-γ labeled nanovials at different cell number to nanovial ratios. The highest single-cell loading efficiency was achieved when cells were seeded at 1.6 cells per nanovial. Increased cell seeding density resulted in a larger fraction of nanovials with two or more T cells inside the cavities of the nanovial. Loading of cells into nanovial cavities followed Poisson statistics. B) Top: Comparison of cell binding motifs (anti-CD45, anti-CD3, anti-CD3/CD28. Nanovials conjugated with anti-CD45 showed the highest loading efficiency when cells were loaded after initial CD3/CD28 activation in culture. Bottom: Increased anti-CD45 concentration on nanovials increased cell binding by nearly 6-fold.C) Flow cytometry analysis and microscopy images showed that nanovials were able to detect each recombinant cytokine (anti-IFN-γ, anti-TNF-α, anti-IL-2) as low as 1 to 10 ng/mL above background with at least three orders of magnitude dynamic range. Each scale bar represents 50 μm.

### Loading of primary T cells onto nanovials

T cells were loaded into the cavities of nanovials by simply pipetting the cell and nanovial suspension in a three-dimensional space such as a well-plate or a tube, followed by incubation for one hour to allow cell binding to anti-CD45 antibodies linked to the nanovial cavities. Single-cell loading onto nanovials through binding with CD45 on the surfaces of T cells was achieved by tuning the cell seeding density to nanovial ratio. When 1.6 cells per nanovial were seeded in a well plate, 15% of nanovials contained single cells and only 3.6% had two or more cells, following expected Poisson loading statistics (Figure 2A). When the cell to nanovial ratio increased, there were more than 10% of nanovials loaded with more than two cells. Depending on the size of the cells, the cell seeding density can be adjusted to ensure that most nanovials have one cell loaded in the cavity. We also tested different types of surface protein targets that can serve as cell binding motifs such as anti-CD3 and anti-CD28, but nanovials labeled with anti-CD45 had the highest loading efficiency because T cells were initially activated for growth with anti-CD3/anti-CD28 soluble activators prior to being loaded, which reduced CD3 and CD28 availability on the cells (Figure 2B). Increased anti-CD45 concentration on nanovials further improved cell binding by nearly 6-fold with nanovials labeled with 210 nM of anti-CD45 having a loading efficiency of 23% compared to a 4% loading efficiency for 70 nM anti-CD45 labeled nanovials (Figure 2B).

### Cytokine detection on nanovials over >3 orders of magnitude dynamic range

We validated the limit of detection of nanovials by pre-labeling nanovials with each cytokine capture antibody and incubating with recombinant proteins (IFN-γ, TNF-α, IL-2). Secreted cytokines were labeled with fluorescent detection antibodies and the signal was analyzed by flow cytometry. Cytokine concentrations as low as 1-10 ng/mL were observable above background with at least three orders of magnitude dynamic range, which we found to be sufficient for single-cell analysis (Figure 2C). When two cytokine capture antibodies were conjugated to the nanovials (against IFN-γ or TNF-α), the ability to detect each individual cytokine was not substantially affected (Figure S1.B). For the same concentration of recombinant IFN-γ or TNF-α, the average intensity on the nanovials only decreased 22% and 31% respectively.

### Measuring single-cell secretion of cytokines on nanovials using FACS

We developed single-cell assays targeting three secreted cytokines (IFN-γ, TNF-α, IL-2) using nanovials coated with separate cytokine capture antibodies and anti-CD45. After loading primary T cells onto nanovials, cells were activated with phorbol 12-myristate 13-acetate (PMA) and ionomycin. Secreted cytokines were accumulated on nanovials over 3 hours and labeled with fluorescent detection antibodies. From fluorescence microscopy images, we observed two distinct fluorescence patterns, fluorescence spread across the nanovial cavity, presumably from secreted cytokines and fluorescence associated with cells on nanovials (without signal on the nanovial). We hypothesized that the fluorescent antibody labels binding to cells could be caused by intracellular staining of permeabilized or dead cells, or by binding to membrane bound forms of cytokines on the cell surface, especially for TNF-α^18^.

We developed an approach to use the fluorescence peak shape to distinguish between nanovial and cell staining. We first looked at the spatial fluorescence distribution in microscopy images (Figure S2.B). From fluorescence images of T cells secreting on nanovials, we plotted the fluorescence intensity profile across the cavity diameter using MATLAB and calculated the maximum intensity (height), the area under the intensity curve (area), and the ratio between the area and height. Nanovials with both spatially spread secretion signal on their cavities and localized labels bound to the surfaces of adhered cells were found to have a similar range of fluorescence peak height values. However, the area over height measurement was distinctly higher for the nanovials with secretion signal. This information may be used as a distinguishing feature in flow cytometry, as analogous fluorescent pulses are generated when nanovials pass through the excitation laser beam spot (Figure S2.A). The height of the flow cytometry pulse is determined by the maximum fluorescence intensity of the nanovial and the area integrates the intensity emitted over the entire transit event through the laser spot. Accordingly, our nanovials with spatially-distributed secretion signals are expected to produce higher fluorescence area signals for a given fluorescence intensity (height) compared to nanovials with cells bound to labels. We analyzed our samples based on a combination of fluorescence peak area and peak height signals (area vs. height plot) and observed two populations, where one population had higher area signal as compared to the other population with similar height values. When we sorted nanovials with larger ratios of area/height (2.06% high secretion and 2.77% low secretion gates), the recovered nanovials had higher secretion signals, while sorted events in the lower area/height (A/H) region (2.61% label binding to cell gate) corresponded to nanovials with label bound to cells (Figure 3A). Using this area vs. height metric, we were able to sort populations of cells with secreted cytokine signal only (A/H > 3), completely differentiating secretion signal on nanovials from signal solely from cell surface binding or intracellular staining of permeabilized or dead cells (Figure 3B). The percent of the cell population with label bound was consistent across all three cytokines (∼2.6% of the total analyzed events). We performed additional quantitative analysis on how the fluorescence peak shape would change with nanovial staining compared to cell staining or a combination of both (see Supplementary Note 1), which provides a guide for differentiating signals from different sources on nanovials and as a function of different sized nanovials.

**Figure 3.**
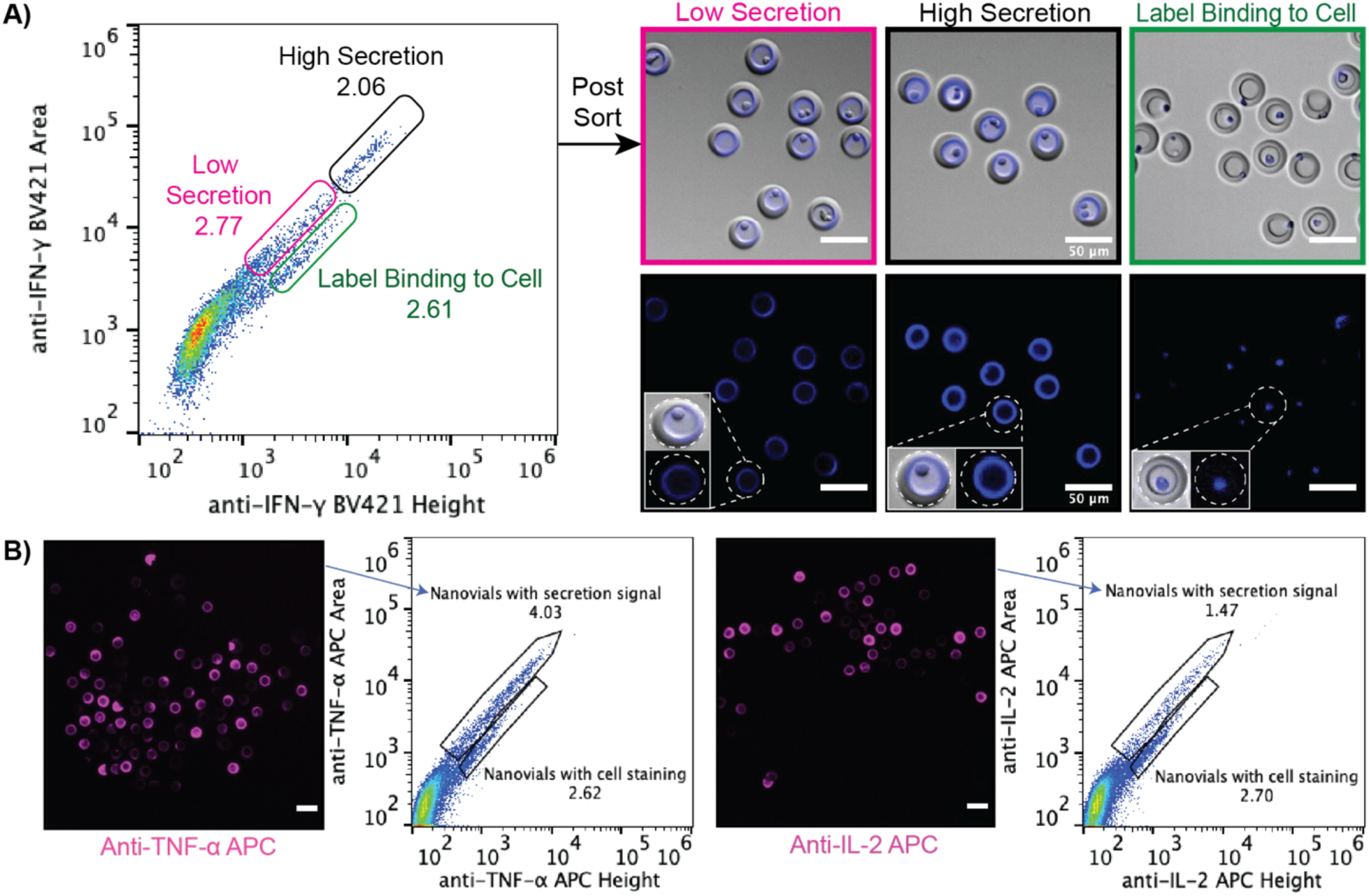
Analysis and sorting of single cells based on cytokine secretion using FACS. A) T cells stimulated with PMA/ionomycin were allowed to secrete for 3 hours on nanovials and analyzed using a Sony SH800. Fluorescence peak area and height are shown where three different gates are used to differentiate spatially-extended IFN-γ secretion signals on nanovials (pink and black secretion gates) from signal solely from non-specific cell surface binding (green label binding to cell gate). Nanovials and loaded cells from these regions were sorted and confirmed to have different distribution of fluorescence signal, as shown in the images. Scale bars represent 50 μm. B) Nanovials with each cytokine (TNF-α or IL-2) secretion signal were sorted using area vs. height metrics. The ability to isolate on-nanovial cytokine staining was consistent across different cytokines as shown in fluorescence microscopy images. Scale bars represent 50 μm.

### Sorting viable cells based on secretion level

We were able to simultaneously measure secretions and viability of individual cells on nanovials, improving the selective sorting of functional cells. Nanovial loaded primary T cells were activated with PMA and ionomycin and secretions were accumulated for three hours. Cells were stained with calcein AM while secreted cytokines were labeled with fluorescently labeled detection antibodies. Using FACS, we first gated our nanovial population based on high calcein AM signal (Figure S2.C) and quantified fluorescence secretion signal using a 2D plot of the fluorescence peak area vs. fluorescence peak height (Figure 4A). We noted that when gating on viable cells the number of events with a low area to height ratio was dramatically reduced, suggesting that fluorescent labeling of cells most likely results from intracellular staining of permeabilized or dead cells. The fluorescent secretion signals associated with viable cells for TNF-α and IFN-γ spanned an order of magnitude in both fluorescence peak area and height. Single cells generally secreting IFN-γ had a distribution that was shifted to higher intensities. Cells secreting TNF-α often had lower fluorescence signal, with a distribution skewed to lower levels.

**Figure 4.**
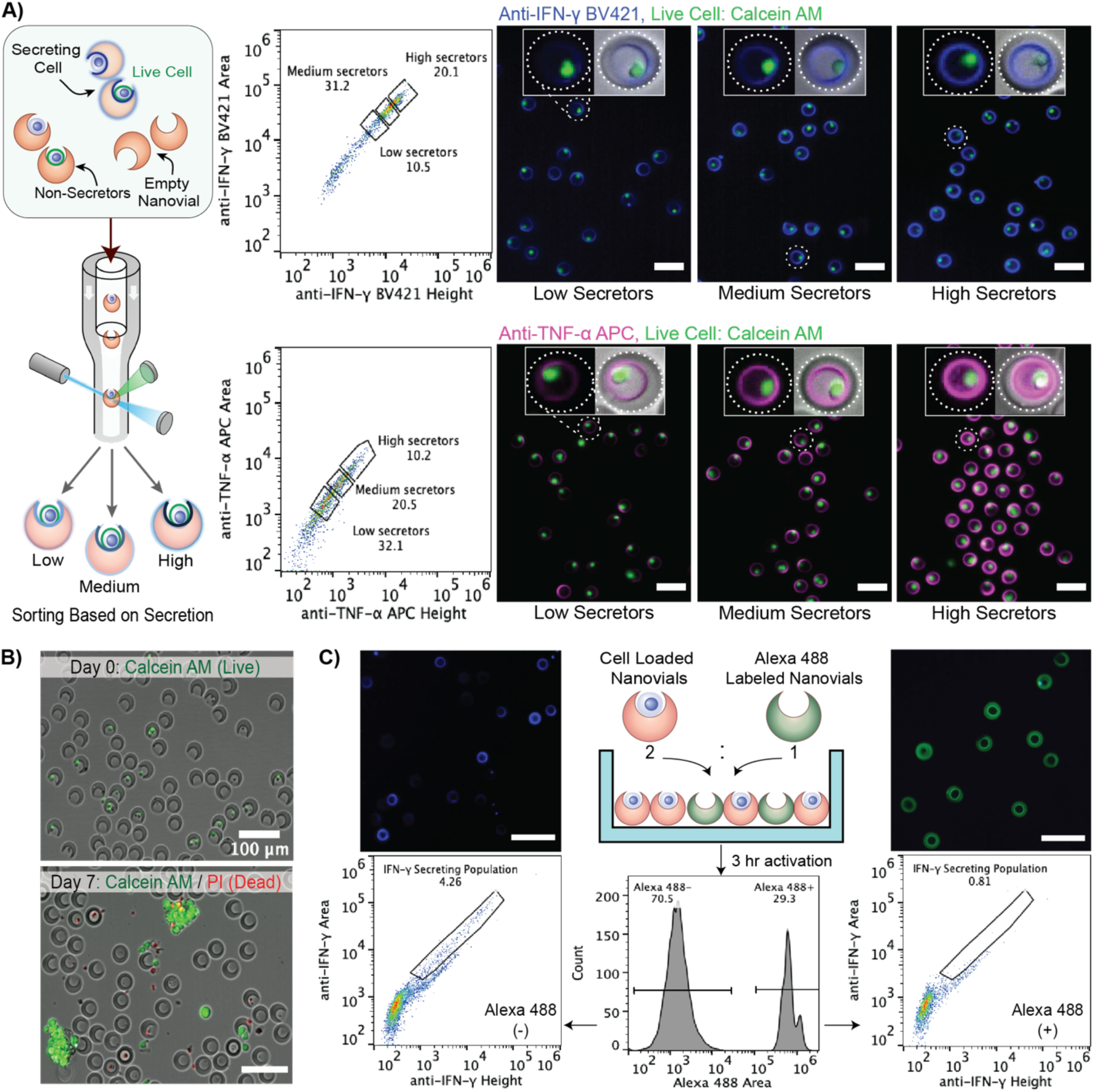
Selection of viable T cells based on cytokine secretion level. A) Captured cytokines and loaded T cells were labeled with fluorescent detection antibodies (anti-IFN-γ BV421 or anti-TNF-α APC) and calcein AM live cell dye. Using a Sony SH800, the nanovial population was first gated for high calcein AM signal (Figure S2.C) and secretion signal was quantified using the fluorescence peak area vs. fluorescence peak height (A/H). Single cells were sorted as high, medium and low secretors based on their IFN-γ and TNF-α secretion level. Microscopy images show each corresponding population, with dotted lines in insets outlining the nanovial boundaries. Scale bars represent 50 μm. B) Recovery and regrowth of cells in nanovials post-sort is seen in microscopy images. Cells increased in number after 7 days, forming in clusters. C) Two nanovial types were introduced together to evaluate cross-talk. Fluorescently labeled nanovials (Alexa Fluor 488) without cells were mixed will T cell-loaded nanovials activated with PMA/ionomycin at a ratio of 1:2. Secretion signal was labeled and analyzed on both nanovial types by gating on the green fluorescence signal on nanovials (Alexa 488 Area). While cell-containing nanovials had significant IFN-γ secretion signal, less than 1% of nanovials without cells had signal in the gate, most at low fluorescence values. Scale bars represent 100 μm.

Viable cells with different levels of TNF-α and IFN-γ secretion could be recovered following sorting. We sorted by first gating on viable cells and then the associated secretion level for TNF-α and IFN-γ. We identified three regimes in the area vs. height plot to gate populations into low, medium, or high secretors. After sorting, cells remained intact and viable in the nanovial, enabling the regrowth of cells post-sort for further downstream analysis. As shown in Figure 4B, cells proliferated over 7 days and formed clusters that grew from nanovials as is expected for T cell culture.

In addition, we incubated cell-loaded nanovials and fluorescently (Alexa Fluor 488) labeled nanovials without cells to determine the crosstalk between two samples. We analyzed secretion signal by gating on the green fluorescence signal on nanovials and found less than 1% of gated nanovials without cells also appeared in our positive secretion gate during the 3 hour activation period (Figure 4C), suggesting minimal cross-talk. Previously, we have seen limited crosstalk in gelatin-functionalized nanovials where the cavity is selectively conjugated with capture antibodies^16,17^. Presumably, secreted cytokines accumulate at higher concentrations locally and are captured but when cytokines diffuse or are advected to neighboring cavities the concentration is diluted substantially leading to a reduced signal.

### Sorting cells based on a combination of secreted cytokines

By decorating the cavity of nanovials with two cytokine capture antibodies, we performed multiplexed secretion-based profiling of >1 million cells to select polyfunctional primary T cells secreting at least two cytokines simultaneously. Nanovials were pre-labeled with two cytokine capture antibodies against IFN-γ and TNF-α along with anti-CD45 antibodies to capture cells. After seeding of cells on the nanovials, cells were activated with PMA and ionomycin and secretions were accumulated over 3 hours. Staining with calcein AM was used to gate out the live cell population only. We first analyzed the secretion signal from nanovials labeled without any cytokine capture antibody and nanovials with each cytokine capture antibody alone to create a positive threshold gate for IFN-γ, TNF-α or IFN-γ and TNF-α secreting cells (Figure S3A). We also stained single-cytokine capture control samples with each detection antibody individually or with both detection antibodies to determine if there was any crosstalk from the detection antibodies. We found individual cytokine signals to be independent of the presence of one or more detection antibodies (Figure S3A).

After we confirmed the accuracy of the multiplexed system and identified positive secretion gates, we performed sorting of T cell populations based on secretion phenotype. We observed that more than half of T cells secreted only IFN-γ, while 4.2% secreted only TNF-α, and 13.4% secreted both cytokines, and were considered polyfunctional T cells. We successfully sorted and recovered the fraction of polyfunctional T cells, which were observed to have staining of the nanovial cavity surfaces with both fluorescent anti-cytokine antibodies (Figure 3B).

### Linking secretions with surface markers from single T cells

We next wanted to identify how the secretion phenotypes identified with the assay correlated with surface markers associated with T cell sub-populations. T cells were loaded onto nanovials conjugated with antibodies against IFN-γ and TNF-α, or antibodies against IFN-γ and IL-2 and anti-CD45 (Figure 5A). After 3 hours of activation with PMA and ionomycin, cells were stained with fluorescent anti-CD4 and anti-CD8 antibodies and captured cytokines were stained with their respective fluorescent antibodies. Our T cell population originally consisted of more CD8+ cells (61.1%) than CD4+ cells (23.5%). When we loaded T cells, 15.9% of nanovials had CD8+ cells while 2.43% of nanovials contained CD4+ cells (Figure 5B). We identified secreting cells by first creating a gate based on the negative control sample that comprised nanovials with only anti-CD45 antibodies. We sorted populations of cells based on fluorescence peak areas exceeding the negative control for each individual cytokine as well as combinations of IFN-γ and TNF-α or IL-2 and IFN-γ (Figure 5C). Cells on nanovials that fell within the negative control gate were considered non-secretors. When measuring IFN-γ and TNF-α, we found that the majority of CD8+ cells secreted IFN-γ and only a small fraction of CD8+ cells secreted TNF-α alone (4.3%) (Figure 5D). About 24% of CD8+ cells were polyfunctional, secreting both IFN-γ and TNF-α simultaneously. On the other hand, CD4+ cells had a larger population that secreted both IFN-γ and TNF-α simultaneously (48%) than CD8+ cells. This pattern was consistent when we analyzed for IFN-γ and IL-2 secretion. About 29% of CD4+ cells secreted both cytokines while 17% secreted IFN-γ only and 13% secreted IL-2 only. Few CD8+ cells secreted IL-2 alone (2%) or IL-2 and IFN-γ (5%). Although the polyfunctional populations were only a small fraction of the entire cell population, we were able to sort and recover them with high viability.

**Figure 5.**
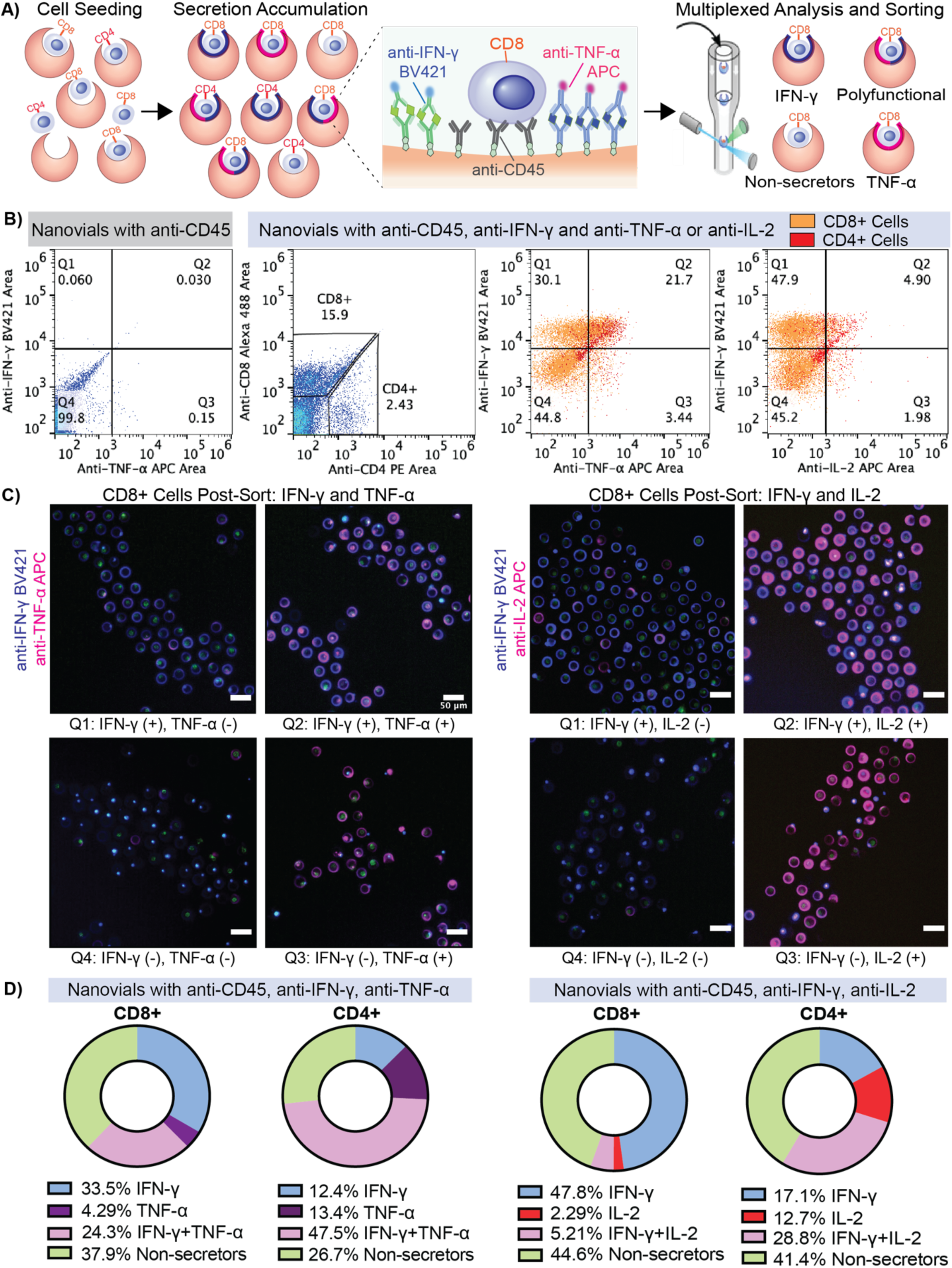
Multiplexed secretion profiling combined with cell surface labeling. A) Overview of multiplexed profiling of single-T cells based on cytokine secretion and cell phenotype. T cells were loaded onto nanovials labeled with anti-IFN-γ and TNF-α, or anti-IFN-γ and IL-2 along with anti-CD45. After accumulating secretion over 3 hours under PMA/ionomycin stimulation, captured cytokines were labeled with their respective fluorescent antibodies and cells were stained with fluorescent anti-CD4 and anti-CD8 antibodies, followed by analysis and sorting with a Sony SH800. B) Flow cytometry dot plots showing CD8+ and CD4+ gates for cells on nanovials, control nanovial dot plots for secretions without any secretion capture antibodies, and experimental dot plots showing IFN-γ and TNF-α or IFN-γ and IL-2 secretion on nanovials. The control nanovial dot plots were used to set the gates for sorting secreting populations of cells on nanovials. C) Single cells were sorted based on CD4+ or CD8+ gates as well as the four quadrant gates shown in B. Scale bars represent 50 μm. D) The population distribution based on secretion is shown as a pie chart for CD4+ and CD8+ cells. There were more polyfunctional CD4+ cells than CD8+ cells.

## Discussion/Conclusion

Nanovials provide a new tool to analyze and sort live T cells based on combinations of secretions and surface markers. This approach has a number of advantages over other currently available single-cell secretion analysis platforms. Nanovials are more accessible than other microfluidic techniques because they can be used with standard FACS machines. Sony SH800 benchtop cell sorter with a 130 micrometer chip was used in this study, and other FACS systems (i.e. BD FACS Aria) are also proven to be compatible with nanovials^19^. Using the Sony SH800 we could sort 350 events per second at sample pressure 4 with over 90% recovery, requiring less than 1 hour to sort 1 million nanovials. All other secretion assay steps can be completed with general lab supplies such as pipettes, tubes, and centrifuges. Other microfluidic approaches to analyze secretions, such as the Berkeley Lights Beacon system^15,20^ or Isoplexis Isolight system^8,21^ require a specialized instrument, which are not as widely available as FACS. We also demonstrate the ability to perform multiplexed surface and secretion-based sorting with <1% crosstalk, leveraging the localized cytokine capture sites on the nanovials. Other reports have found that the Miltenyi cytokine catch assay can result in significant crosstalk to neighboring capture-antibody labeled cells^14^, and a reduced dynamic range since cytokines can be easily transported away from secreting cells. Also, the crosstalk and dynamic range of nanovials can be further improved by encapsulation of nanovials in oil to highly concentrate secreted cytokines while maintaining the viability of cells^16^. Another key differentiator from other techniques is that nanovials can recover viable cells for further downstream analysis. This is not possible with ELISpot, ICS or the Isolight.

Our nanovial technology can be easily scaled up to process millions of cells per day. Here, we fabricated nanovials using a flow-focusing device at rate of 5.6 million/hour, and higher production rates can be achieved using parallelized microfluidic device^17^. For each secretion assay, we loaded 0.23 million cells into 187,000 nanovials in a 24-well plate yielding ∼28,000 cells loaded in nanovials. Among those bound, 20-30% of cells secreted cytokines and were sorted. By using larger containers (e.g. a 10 cm Petri dish or centrifuge tubes) for loading and increasing the number of cells introduced, we can prepare and analyze millions of cells. Recently, we were able to scale up our secretion assay to perform cytokine secretion-based sorting of a starting population of 10 million T cells loaded into 3 million nanovials all within a 3 hour sorting period (data not shown). Here the bottleneck for throughput becomes the flow cytometer event rate^22^. Other microfluidic platforms have been limited to a few thousand cells^23^ or a maximum of 50,000 cells per run^15,20^. The current throughput can be further increased by reducing the size of the nanovials and decreasing the nozzle size of the FACS machine to process nanovials at a rate that could approach 10,000 events per second.

Using a combination of fluorescence peak area and height signals in FACS, we have shown that we can differentiate secreted cytokines captured on the nanovial cavity from other fluorescence labeling likely due to intracellular staining of permeabilized or dead cells. When using viability dye, this cell labeling could be completely gated out. This analytical feature is particularly important for some secreted proteins as they can also be present on the cell surface. For example, there are both secreted and soluble forms of TNF-α, which would reflect different biological states of a cell. TNF-α can be expressed on the surface of activated T cells where it exerts various pro-inflammatory functions in a cell-to-cell contact manner, which are distinct from the soluble form of TNF-α. Membrane TNF-α has been shown to be significantly induced on the cell surface of CD8+ T cells from autoimmune disease patients where it exerts cytotoxic activity.^18^. Similarly, for B cells, secreted IgG and the B cell receptors share a common structure^24^. Therefore, the area/height ratio, or other metrics of fluorescence peak width or shape, can be used to distinguish these different states, unlike other approaches like ICS or the Miltenyi cytokine catch assay in which all staining occurs on the cell surface or intracellularly and cannot be distinguished^13,25–28^. Images from image cytometry systems^29,30^ or image-activated cell sorting systems^31,32^ would provide the similar ability to distinguish between cell surface staining and staining on nanovials.

By staining T cells with additional CD4 and CD8 markers, we were able to relate secretion characteristics with corresponding cell phenotype and compare our results to previous studies using different approaches to measure single-cell secretions. Among T cell populations, we showed that the majority of CD8+ cells secreted IFN-γ upon PMA and ionomycin activation and this finding is in line with a previous study, using a microengraving based secretion assay, which reported 30% of CD8+ cells secreted IFN-γ after 6 hours of PMA and ionomycin activation and a limited population of TNF-α (5%) and IL-2 (2%) secreting cells^23^. A microarray approach reported no IL-2 after 6 hours of PMA and ionomycin activation and an ICS-based study reported 39% of CD8+ were IFN-γ positive and only 2% were IL-2 positive^33^. However, the polyfunctional population of CD8+ cells we reported was significantly higher than what was found in the previous microengraving study that detected only 1% IFN-γ and TNF-α and 2% IFN-γ and IL-2 secreting populations. Similarly for CD4+ T cells, we found a large population of polyfunctional T cells but previous microengraving findings indicated only a small fraction of cells secreted multiple cytokines (8% IFN-γ and IL-2, 1% IFN-γ and TNF-α)^23^. Our CD4+ cell secretion characteristics were also different from other ICS studies which discovered a much higher number of cells secreting each cytokine (30-50% IFN-γ, 47% TNF-α, or 45% IL-2) after 6 hours stimulation with PMA and ionomycin^33–35^.

These discrepancies may arise from differences between the studies or readout methods. In our studies, T cells are under proliferative culture conditions (CD3/CD28 stimulation) prior to analysis, which may lead to differences in the accumulation of specific cytokines prior to stimulation. We also looked at an early time point. Because of the higher sensitivity of the nanovials to secreted cytokines we evaluated cytokines accumulated up to 3 hours after stimulation in our studies vs. 6 hours in other studies. There are also slight differences in PMA/ionomycin stimulation concentrations from study to study. We may not expect direct correlations with ICS, which only allows staining of accumulated cytokines that are intracellular and may not represent the actual amount of cytokines that are secreted. Given our larger sample sizes in which we analyzed a total of 24,000 CD8+ and CD4+ cell loaded nanovials as compared to 3,015 CD8+ and CD8-cells in the microengraving study, we can be more confident in the statistics of the populations, even for rare populations of secreting cells. Our results with CD4+ cells may be affected by previous reports that indicate downregulated CD4 expression in response to PMA/ionomycin stimulation, although we expect this to be less of an effect for the shorter 3 hour time point we use in our studies.

This is the first application of nanovials to sorting of cells based on more than one secreted product as well as combined sorting based on secreted and surface markers, opening up new applications in immunology and cell therapy discovery. Previous work has identified that the presence of polyfunctional T cells which secrete more than one cytokine are associated with improved responses in patients undergoing CAR-T cell therapy. In this study, we have specifically analyzed three dominant cytokines (IFN-γ, TNF-γ and IL-2) secreted by T cells upon PMA and ionomycin activation but our nanovials can be further decorated with other functional groups such as antibodies against effector molecules (granzyme B, MIP-1a, perforin), regulatory cytokines (IL-10, IL-13 and IL-17) and inflammatory cytokines (IL-6, MCP-1). Another key benefit of our technology is that sorted cells can be recovered and investigated further for downstream function (e.g. with *in vitro* or animal models) or transcriptomic information (either pooled or using single-cell sequencing technologies). In our previous studies, we have shown that the morphology of the nanovial cavity protects cells during damaging sorting steps, leading to improved viability, cell expansion and proliferation post-sort^16,17^. The ability to functionally sort viable intact cells will allow improved understanding of unique polyfunctional phenotypes. Future application of nanovials include high-throughput sorting of rare antigen-specific T cells based on secreted cytokines that are accumulated on nanovials. Similarly, CAR-T cells can also be selected and sorted based on binding to antigen-labeled nanovials and secretion of cytokines indicating functional engagement. From sorted polyfunctional T cells, antigen-specific T cells, and CAR-T cells it is possible to perform transcriptomic analysis to uncover molecular drivers of this phenotype and sequence TCRs or CARs from the enriched population to find novel functional engineered TCR constructs or CARs from a library leading to optimal functional responses for therapeutic use.

## Methods

### Nanovial fabrication

Nanovials were fabricated using a three-inlet flow-focusing microfluidic droplet generator formed from polydimethylsiloxane as previously described^16^. In respective inlets, PEG pre-polymer, gelatin and oil phases were infused at flow rates of 1.5 μl/min, 1.5 μl/min, and 15 μl/min respectively. The PEG pre-polymer phase comprised 27.5% w/v 5kDa 4-arm PEG acrylate (Advanced BioChemicals) with 4% w/v lithium phenyl-2,4,6-trimethylbenzoylphosphinate (LAP, Sigma) in phosphate buffered saline (PBS, pH 7.2). The gelatin phase comprised 20% w/v cold water fish gelatin (Sigma) in deionized water. The oil phase comprised Novec 7500 (3M) with 0.5% v/v Pico-surf (Sphere Fluidics). Oil partitioned the aqueous phases into monodisperse water-in-oil droplets and PEG-gelatin polymers phase separated after approximately 5 seconds, followed by cross-linking with focused UV light through a DAPI filter set and microscope objective (Nikon, Eclipse Ti-S) near the outlet region of the microfluidic device. Polymerized nanovials were collected in a conical tube and any unreacted phases including oil were removed through a series of washing steps as previously described^16,17^. Biotinylation of the gelatin-layer formed in the nanovial cavity was conducted by incubating nanovials with 10 mM Sulfo-NHS-Biotin (APExBIO) overnight at 4°C. Nanovials were then washed in Washing Buffer consisting of 0.05% Pluronic F-127 (Sigma), 1% 1X antibiotic-antimycotic (Thermo Fisher), and 0.5% bovine serum albumin (BSA, Sigma) in PBS and sterilized in 70% ethanol overnight. Sterile nanovials were stored at 4°C in Washing Buffer.

### Antibody conjugation to nanovials

Sterile nanovials were diluted in Washing Buffer five times the volume of nanovials (i.e. 100 μL of nanovial volume was resuspended in 400 μL Washing Buffer). Diluted nanovial suspension was incubated with equal volume of 200 μg/mL of streptavidin for 30 minutes at room temperature on a tube rotator. Excess streptavidin was washed out three times by pelleting nanovials at 200g for 5 minutes, removing supernatant and adding 1 mL of fresh Washing Buffer. Streptavidin-coated nanovials were reconstituted at a five time dilution in Washing Buffer containing 140 nM (20 μg/mL) of each biotinylated antibody or cocktail of antibodies: anti-CD45 (Biolegend) and anti-IFN-γ (R&D Systems), anti-TNF-α (R&D Systems), or anti-IL-2 (BD Sciences). Nanovials were incubated with antibodies for 30 minutes at room temperature on a rotator and washed three times as described above. Nanovials were resuspended at a five times dilution in Washing Buffer or culture medium prior to each experiment.

### T cell culture

T cells were isolated from human donor whole blood samples by negative selection using the RosetteSep Human T Cell Enrichment Cocktail Kit (STEMCELL Technologies). Isolated T cells were seeded in a fresh complete ImmunoCult™-XF T cell expansion medium (STEMCELL Technologies) at 1 × 10^6^ cells/mL with 2 μL/mL ImmunoCult™ human CD3/CD28 T cell activator. T cells were activated for 3 days and expanded for up to 12 days by changing into fresh expansion medium every 2-3 days. All cells were cultured in incubators at 37°C and 5% CO_2_.

### Cell loading into nanovials

T cells were stained with 1 μM cell tracker deep red dye (Thermo Fisher) for 30 minutes and loaded into nanovials as previously described, with some modification^16,17^. Each well of a 24-well plate was filled with 1 mL of T cell expansion medium and 30 μL of reconstituted anti-CD45 antibody-labeled nanovials (6 μL of nanovial volume=187,000 total nanovials) were added in each well using a standard micropipette. Cells were seeded in each well and extra culture medium was added to make a total volume of 1.5 mL. Each well was mixed by simply pipetting 5 times with a 1000 μL pipette set to 1000 μL. The well plate was transferred to an incubator to allow cell binding for one hour; each well was pipetted up and down again with a 200 μL pipette at 30-minute intervals. After one hour, nanovials were strained to remove any unbound cells and recovered using a 20 μm cell strainer. During this step, any unbound cells were washed through the strainer and only the nanovials (with or without cells loaded) were recovered into a conical tube or well plate.

### Cell loading efficiency and statistics

Nanovials labeled with anti-CD45 antibodies were prepared using the procedures described above. To test concentration dependent loading of nanovials 0.15 × 10^6^ (0.8 cells per nanovial), 0.3 × 10^6^ (1.6 cells per nanovial), and 0.47 × 10^6^ (2.4 cells per nanovial) of cell tracker deep red stained T cells were seeded in each well of a 24-well plate along with 187,000 nanovials. After incubating for one hour in the incubator to promote cell binding, nanovials were recovered and transferred to a well plate to be imaged with a fluorescence microscope. Loading efficiency was analyzed using a custom image analysis algorithm in MATLAB which detected nanovials using standard segmentation algorithms. The software measured the total number of nanovials in each image frame, then the number of cells in each nanovial was manually counted to record the total number of nanovials with 0, 1 or 2 or more cells (n > 2000). For comparing different cell binding motifs nanovials were labeled with 140 nM of each biotinylated antibodies (anti-CD3, anti-CD3 and anti-CD28, or anti-CD45) and seeded with 0.3 million cells in each well. To determine the effect of increased anti-CD45 concentration on nanovials, nanovials were labeled with 0, 70, 140, or 210 nM of anti-CD45 antibodies and 0.3 million cells were seeded in each well with nanovials in a 24-well plate. After cell binding and recovery of nanovials, the number of cells in each nanovial were analyzed using the same image analysis algorithms mentioned above (n > 2000).

### Flow cytometer analysis and sorting

All flow cytometry analysis and sorting were performed using the SONY SH800 cell sorter equipped with a 130 micron sorting chip (Catalog: LEC3213). The cytometer was configured with violet (405 nm), blue (488 nm), and red (640 nm) lasers with 450/50 nm, 525/50 nm, and 665/30 nm filters. Standard gain settings for different sensors are indicated in table 1 below. In each analysis, samples were compensated using negative (blank nanovials) and positive controls (1000 ng/mL recombinant cytokine captured nanovials labeled with each fluorescent detection antibodies). Nanovial samples were diluted to approximately 0.062 million per mL in washing buffer for analysis and sorting. Drop delay was configured using standard calibration workflows and single-cell sorting mode was used for all sorting as was previously determined to achieve the highest purity and recovery^19^. A sample pressure of 4 was targeted.

**Table 1.** Common gain settings used for analysis and sorting.

| Sensor | FSC | BSC | FL1 | FL2 | FL4 |
| --- | --- | --- | --- | --- | --- |
| Gain | 1 | 26% | 28% | 22% | 30% |

### Dynamic range of cytokine detection on nanovials

Nanovials were labeled with biotinylated antibodies (140 nM anti-CD45 and 140 nM anti-IFN-γ, anti-TNF-α or anti-IL-2) using the modification steps mentioned above. Each sample of cytokine capture antibody-labeled nanovials was incubated with 0, 10, 100, or 1000 ng/mL of recombinant human IFN-γ (R&D Systems), TNF-α (R&D Systems), and IL-2 (R&D Systems) for 2 hours at 37°C. Excess proteins were removed by washing nanovials three times with washing buffer. Nanovials were pelleted at the last wash step and incubated with 5 μl of 100 μg/mL anti-IFN-γ BV421 (Biolegend, Catalog: 502532), 25 μg/mL anti-TNF-α Alexa Fluor 647 (Biolegend, Catalog: 502916) and 50 μg/mL anti-IL-2 Alexa Fluor 488 (Biolegend, Catalog: 500314) in 50μL washing buffer per 6 μL nanovial volume at 37°C for 30 minutes protected from light. Following washing three times, nanovials were reconstituted at a 50 times dilution in the washing Buffer and transferred to a flow tube. A small fraction of the sample was transferred to a 96-well plate to be imaged on a fluorescence microscope. Fluorescent signal on nanovials was analyzed using a SONY SH800 with sensors and gains mentioned in flow cytometer analysis and sorting section.

### Analyzing single-cell cytokine secretion on nanovials

Nanovials were sequentially coated with streptavidin and biotinylated antibodies (140 nM anti-CD45 and 140 nM anti-IFN-γ, anti-TNF-α or IL-2). In each well of a 24-well plate, antibody labeled nanovials and 0.3 million cells were mixed in a total of 1.5 mL T cell expansion medium to allow cell binding. Any unbound cells were removed by a 20 μm cell strainer and nanovials were recovered into a 12-well plate in 2 mL of T cell expansion medium with 100 ng/mL PMA and 2.5 μM Ionomycin. T cells were activated for 3 hours in a 37°C incubator to accumulate secretion and nanovials were collected in a conical tube with Washing buffer. After 5 minutes of centrifugation at 200g, supernatant was removed and nanovials were reconstituted at a ten-fold dilution in Washing buffer containing 5 μl of 100 μg/mL anti-IFN-γ BV421 (Biolegend, Catalog: 502532), 25 μg/mL anti-TNF-α APC (Biolegend, Catalog: 502912) and 200 μg/mL anti-IL-2 APC (BD Biosciences, Catalog: 554567) per 6 μL nanovial volume. Nanovials were incubated at 37°C for 30 minutes, protected from light to label secreted cytokines. After washing nanovials with 5 mL of washing buffer, nanovials were resuspended at a 50-fold dilution in the washing buffer and transferred to a flow tube, while a small fraction of sample was transferred to 96-well plate to be imaged prior to sorting on a fluorescence microscope. Pre-sort images were analyzed by custom image analysis algorithms in MATLAB. Fluorescence intensity profiles were calculated along a line segment manually defined around the cavity of nanovial. The intensity peak height and the area under the intensity profile were then evaluated, to find the peak area over height aspect ratio. In FACS, samples were analyzed based on a combination of fluorescence area and height signals with sensors and gains mentioned in flow cytometer analysis and sorting section. The population with higher area signal (as compared to the other population with similar height values) was sorted as nanovials with secretion signal, while the other population was sorted as nanovials with intracellular staining. For each cytokine, 40,000 nanovials were analyzed and directly sorted into a 96-well plate and imaged with a fluorescence microscope to quantify the enrichment of nanovials with secretion signal or intracellular staining.

### Sorting of high, medium, and low secretors

Nanovials were labeled with streptavidin and biotinylated antibodies (140 nM anti-CD45 and 140 nM anti-IFN-γ or anti-TNF-α). Antibody-labeled nanovials (187,000) and 0.3 million cells were mixed in a total of 1.5 mL T cell expansion medium to allow cell binding for one hour in a 24-well plate. Nanovials were recovered using a 20 μm cell strainer and seeded into a 12-well plate in 2 mL of T cell expansion medium with 100 ng/mL PMA and 500 ng/mL ionomycin. T cells were activated for 3 hours in a 37°C incubator to accumulate secretion and nanovials were reconstituted at a ten times dilution in washing buffer containing 0.3 μM calcein AM dye (Thermo Fisher) along with 5 μL of each detection antibody (anti-IFN-γ BV421, anti-TNF-α APC, anti-IL-2 APC) per 6 μL nanovial volume. Nanovials were incubated at 37°C for 30 minutes, protected from light. Nanovials were resuspended at a 50 times dilution in the washing buffer and transferred to a flow tube to be analyzed by FACS. To sort live single cells based on secretion signal, nanovials with calcein staining were first gated and high, medium, or low secretors were sorted by thresholding the fluorescence area and height signals as shown in the results. Sorted samples were imaged with a fluorescence microscope to validate the enrichment of nanovials based on the amount of secretion captured on the nanovials.

### Linking cell surface markers to secretion phenotype

We performed analysis of secretions across multiple cytokines and linked this to surface marker expression. Streptavidin-coated nanovials were decorated with biotinylated antibodies (140 nM anti-CD45, 140 nM anti-IFN-γ, 140 nM anti-TNF-α or 140 nM anti-CD45, 140 nM anti-IFN-γ, 140 nM IL-2) for 30 minutes. Negative control nanovials were also prepared by labeling nanovials only with anti-CD45 antibody without any cytokine capture antibodies. In each well of a 24-well plate, nanovials were seeded with 0.5 million cells in a total of 1.5 mL T cell expansion medium. Cells were allowed to bind for 1.5 hour in a 37°C incubator. After straining and recovery of nanovials, cells were activated for 3 hours in the T cell expansion medium containing 100 ng/mL PMA and 500 ng/mL ionomycin. To stain secreted cytokines nanovials were washed once with 5 mL washing buffer and reconstituted at a ten times dilution in washing buffer containing 5 μL of each detection antibody (anti-IFN-γ BV421, anti-TNF-α APC, anti-IL-2 APC), 5 μL of 25 μg/mL anti-CD4 PE (Biolegend, Catalog: 344606) and 5 μL of 100 μg/mL anti-CD8 Alexa Fluor 488 (Biolegend, Catalog: 344716) per 6 μL nanovial volume for 30 minutes in a 37°C incubator. Nanovials were resuspended at a 50 times dilution in the washing buffer and transferred to a flow tube to be analyzed in FACS. In flow, nanovials with CD4 and CD8 cells were gated based on fluorescence area signals of Alexa Fluor 488 and PE. Nanovials with secretion signal were evaluated by first creating quadrant gates based on the negative control sample (nanovials only labeled with anti-CD45 antibody). Q1 was defined as nanovials with only IFN-γ secreting cells. Q2 was nanovials with polyfunctional T cells that secreted both cytokines (IFN-γ and TNF-α or IL-2). Q3 was nanovials with either TNF-α or IL-2 secreting cells while Q4 was nanovials with non-secretors. Nanovials in each quadrant were sorted into a 96-well plate and samples were imaged with a fluorescence microscope to quantify enrichment of each cell type and their associated secretion characteristics. For each multiplexed screening, 100,000 nanovials were analyzed and sorted by FACS per condition and each condition was repeated three times for a total of 300,000 nanovials analyzed per condition.

## Acknowledgements

This work was supported by the National Institutes of Health through award R21 CA256084. Flow cytometry was performed in the UCLA Jonsson Comprehensive Cancer Center (JCCC) and Center for AIDS Research Flow Cytometry Core Facility that is supported by National Institutes of Health awards P30 CA016042 and 5P30 AI028697, and by the JCCC, the UCLA AIDS Institute, the David Geffen School of Medicine at UCLA, the UCLA Chancellor’s Office, and the UCLA Vice Chancellor’s Office of Research. We thank Jamie Spangler and Monika Kizerwetter for helpful discussions.

