## Supplementary Information for "Sorting single T cells based on secreted cytokines and surface markers using hydrogel nanovials"

### Table of Contents

Supplementary Note 1………………………………………………………………… 2
Figure S1………………………………………………………………………….. 6
Figure S2………………………………………………………………………….. 7
Figure S3………………………………………………………………………….. 8

**Supplementary Note 1. Mathematical rationale for distinguishing nanovial specific signal and cell specific signal using fluorescence area to height ratio**

When performing secretion assays we expect fluorescence signal localized to the nanovial. However, microscopy images show that some populations may have intracellular uptake that can lead to misclassification as secreting populations. Experimentally we showed that these populations can be distinguished by looking at the ratio of the area and height of the measured fluorescence pulse using flow cytometry. In general, we find this to a more robust way to differentiate populations then just looking at peak width as some cytometers have a fixed measurement width across all channels (e.g. the SONY SH800 bases width off only the event triggering channel.

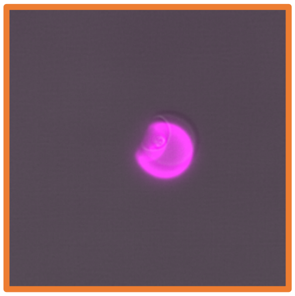
Nanovials with secretion signal

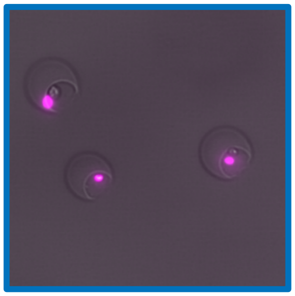
Labels binding to cells

Here we create a simple mathematical model to estimate the expected pulse signal for different staining configurations and support our experimental observations.

To simplify calculations, we will treat both the cells and nanovials as spherical objects and approximate their fluorescence pulse profile as a gaussian distribution:

$$Fluorescence\ Intensity = F(t) = H \exp\left(-\frac{(t-t_0)^2}{2W^2}\right)$$

where H is the pulse height, W is the pulse width, $t$ is time, and $t_0$ is the initial time of the intensity pulse.

We can calculate the area of the pulse by integrating the above expression from $t = -\infty$ to $t = -\infty$ and set $t_0 = 0$:

$$\text{A} = \int F(t) = \int H \exp\left(-\frac{(t)^2}{2W^2}\right) dt = \frac{1}{2}\sqrt{2\pi}H * W \operatorname{erf}\left(\frac{t}{W\sqrt{2}}\right) + const$$

$$\int_{-\infty}^{+\infty} F(t) = \sqrt{2\pi}HW$$

Using this formula, we plotted the pulse profiles for cells and nanovials with different intensities along with the summation of the profiles to predict signal from cells loaded inside of a nanovial. These example fluorescence pulse heights, areas and area/height ratios are given in the table below.

| Sample | Cell | | | | Nanovial | | | | Combined | | | |
|---|---|---|---|---|---|---|---|---|---|---|---|---|
| | H | W | Area | Area/H | H | W | Area | Area/H | H | W | Area | Area/H |
| 1) Bright Cell Dim Nanovial | 1 | 1 | 2.507 | 2.507 | 0.25 | 4 | 2.507 | 10.03 | 1.25 | NA | 5.014 | 4.011 |
| 2) Dim Cell Bright Nanovial | 0.25 | 1 | 0.627 | 2.506 | 1 | 4 | 10.03 | 10.03 | 1.25 | NA | 10.66 | 8.528 |
| 3) Dim Cell Dim Nanovial | 0.25 | 1 | 0.627 | 2.506 | 0.25 | 4 | 2.507 | 10.03 | 0.5 | NA | 3.134 | 6.268 |
| 4) Bright Cell Bright Nanovial | 1 | 1 | 2.507 | 2.507 | 1 | 4 | 10.03 | 10.03 | 2 | NA | 12.54 | 6.270 |

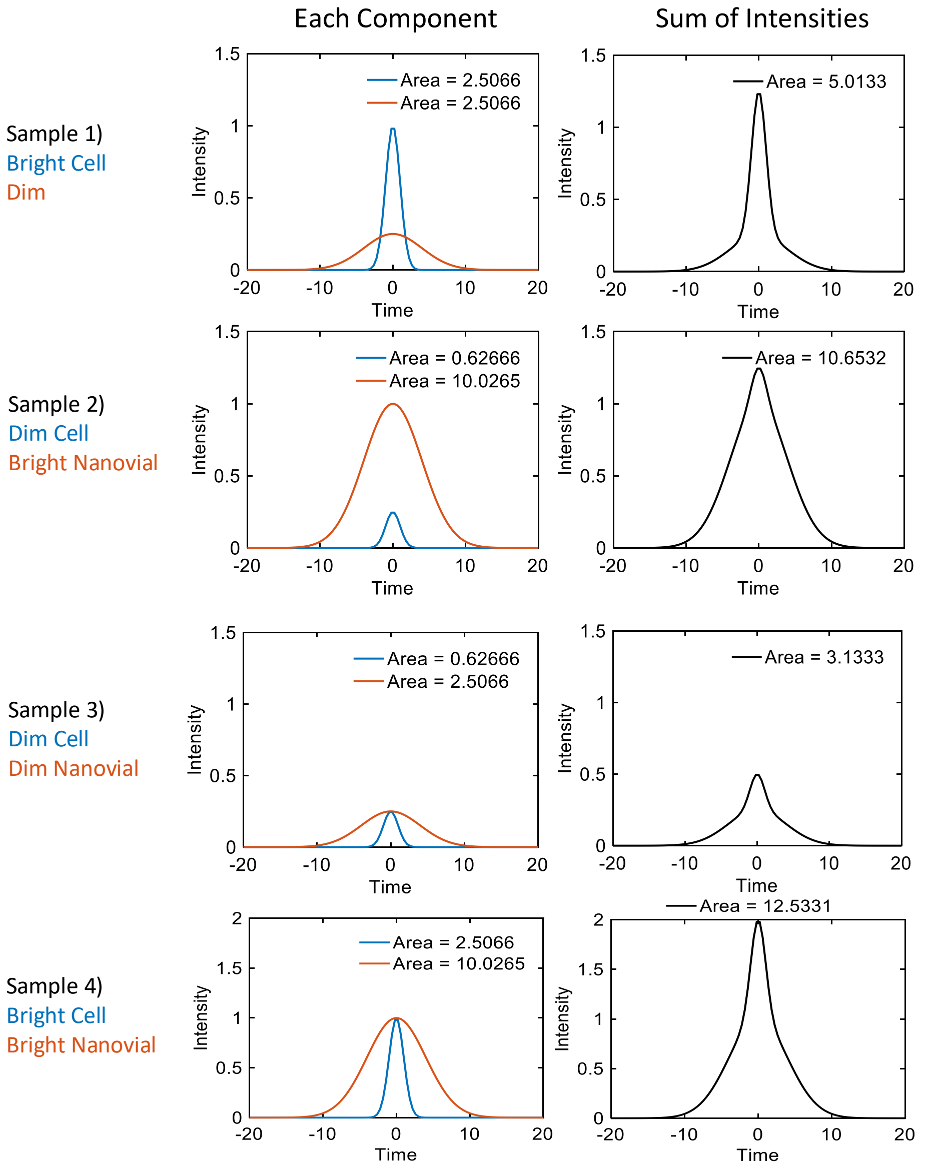

Mathematically the pulse area to pulse height ratio for the combined pulses of the cell ($F_C$) and nanovial ($F_N$), can be expressed as:

$$\frac{A_{CN}}{H_{CN}} = \frac{\sqrt{2\pi}(H_C W_C + H_N W_N)}{H_C + H_N}$$

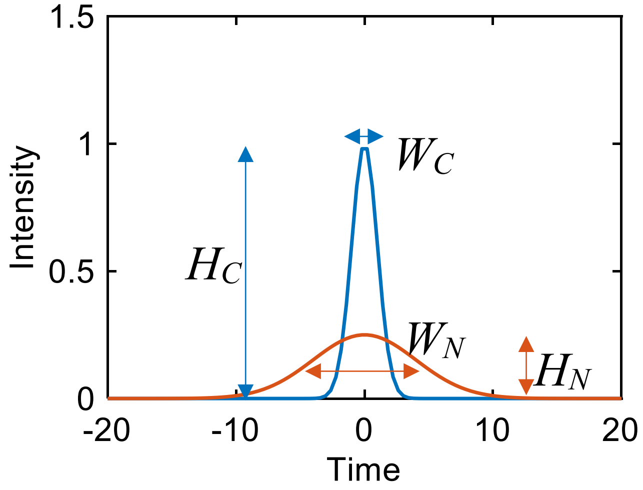


If we then take this ratio and plot it as a function of the pulse height ratio we see that as the height of the nanovial fluorescence pulse increases relative to the fluorescence pulse of the cell the area over height value increases and asymptotically approaches a value that is dependent on the nanovial width. Likewise, as the cell fluorescence pulse dominates, we see that the value asymptotically approaches a value that depends on the width of the cell pulse.

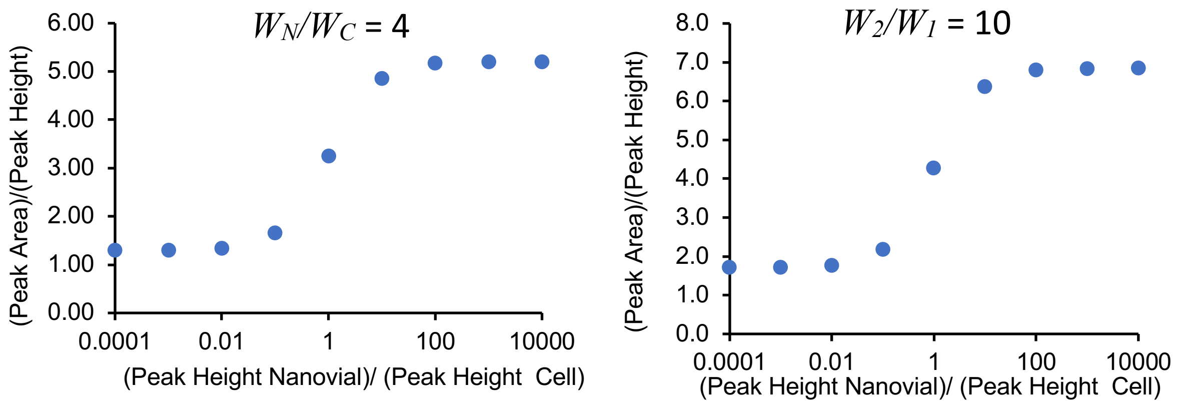

Using FACS we can measure both the Peak Area and Peak Height. If we plot a dot plot of nanovial events as a function of fluorescence height and fluorescence area, nanovial dominated fluorescence events will fall on a higher slope line (offset line to the top left in the log-log plot). Therefore, by gating populations with high A/H ratios we can analyze and sort nanovials and attached cells with nanovial staining associated with secretion and not cell surface or intracellular staining.

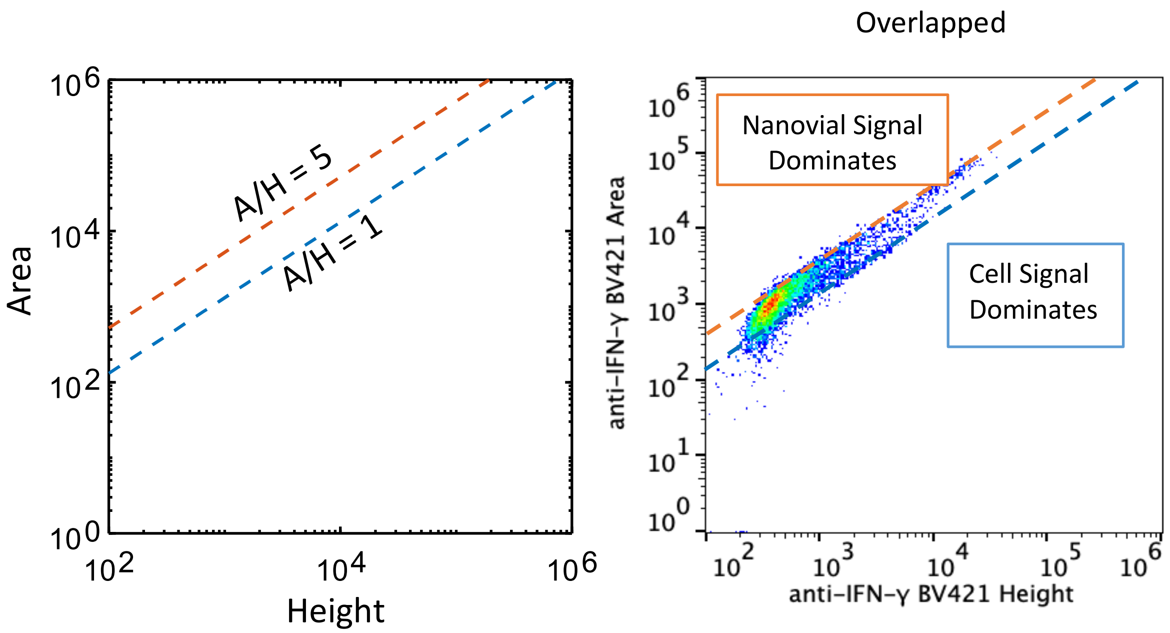

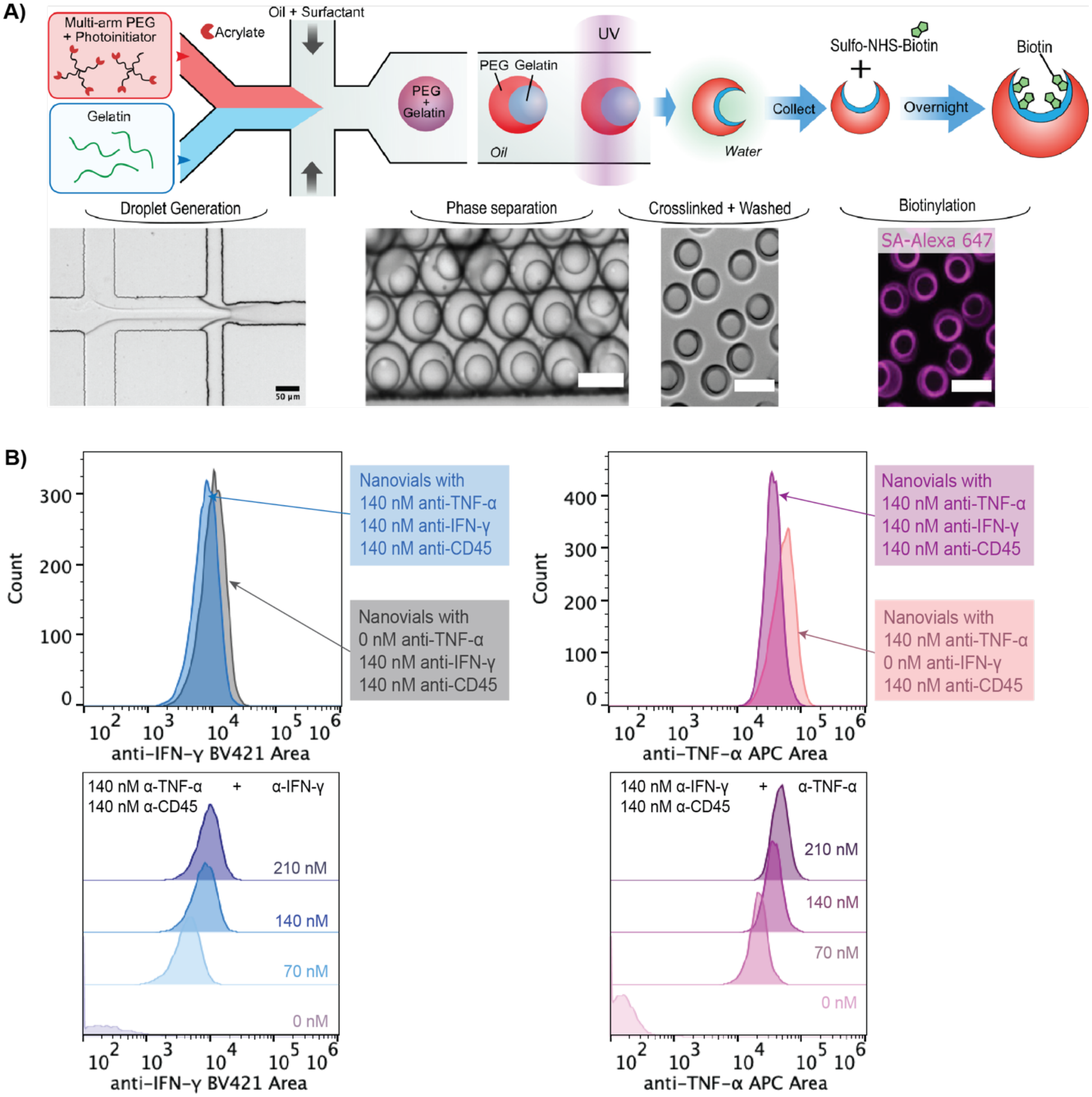


Figure S1 Nanovial fabrication and functionalization with multiple antibodies. A) An aqueous phase comprised of 4-arm PEG acrylate and photoinitiator is co-flowed with a gelatin solution in a microfluidic droplet generator to make uniform monodisperse aqueous two-phase water-in-oil droplets. After droplet formation, PEG and gelatin undergo phase separation and then are exposed to UV light at the end of the device to cross-link the 4-arm PEG acrylate. After collection, any excess polymerized PEG, gelatin, or oil are removed during washing and cross-linked nanovials are incubated with Sulfo-NHS-biotin to be biotinylated. Scale bars represent 50 µm. B) Flow cytometry fluorescence histograms of nanovials following the cytokine capture assay with different capture antibody concentrations and recombinant cytokines. Nanovials were functionalized with cell binding motif (anti-CD45) and two cytokine capture antibodies at different concentrations (anti-IFN-γ, anti-TNF-α) and were incubated with recombinant TNF-α or IFN-γ. The ability of nanovials to detect each individual cytokine was not significantly affected with the presence of other cytokine capture antibody. There was also a capture antibody concentration-dependent shift in the average intensity on the nanovials.

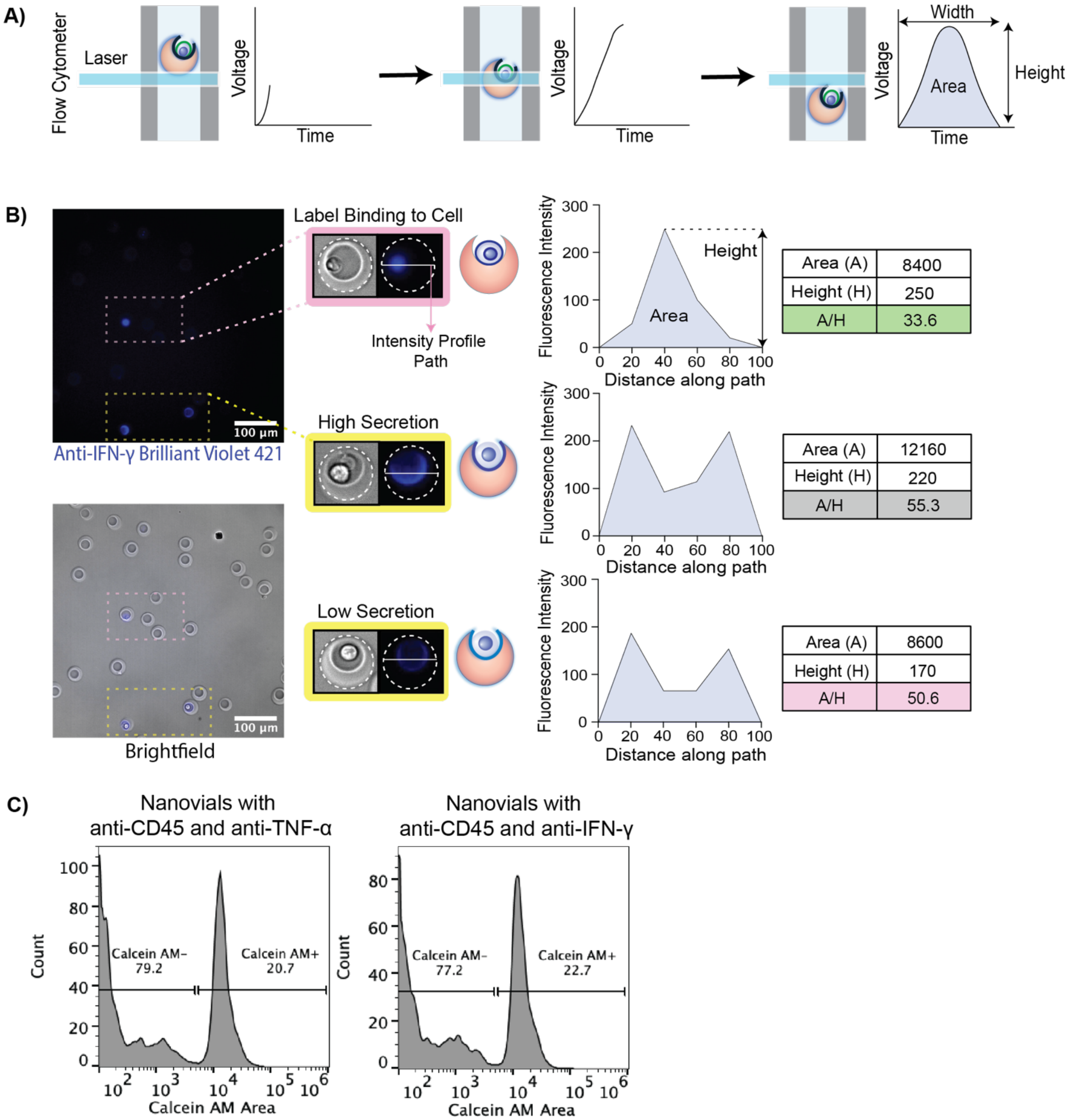


Figure S2 Using the fluorescence peak area and height to measure single-cell secretions on nanovials. A) Overview schematic of the nanovial fluorescence peak shape obtained when a nanovial transits a laser spot in a flow cytometer. A nanovial passes through the laser beam and generates scattered light and fluorescence signals in a time-dependent manner. The nanovial is fully illuminated and produces a maximum amount of fluorescence signal (Peak Height) when it is located at the center of the laser and as it flows out the signal drops back to the baseline. The area is the integral of the fluorescence intensity emitted over the entire transit event through the laser spot. B) Fluorescence microscopy images of pre-sort nanovials with cells showed two distinct fluorescence patterns, with dotted lines in insets outlining the nanovial boundaries: fluorescence spread across the nanovial cavity from secreted cytokines or fluorescence associated with cells on nanovials without signal across the cavity area. The fluorescence intensity profile was computed across the cavity of each image and the maximum intensity (height, H), area under the intensity curve (area, A), and the ratio between the area and height were calculated. Nanovials with spatially spread secretion signal and nanovials with label bound to cells had similar peak height values, but secretion signal had much higher area over height ratio (A/H). Scale bars represent 100 µm. C) Nanovials were first gated for high calcein AM signal to select viable cells for secretion analysis.

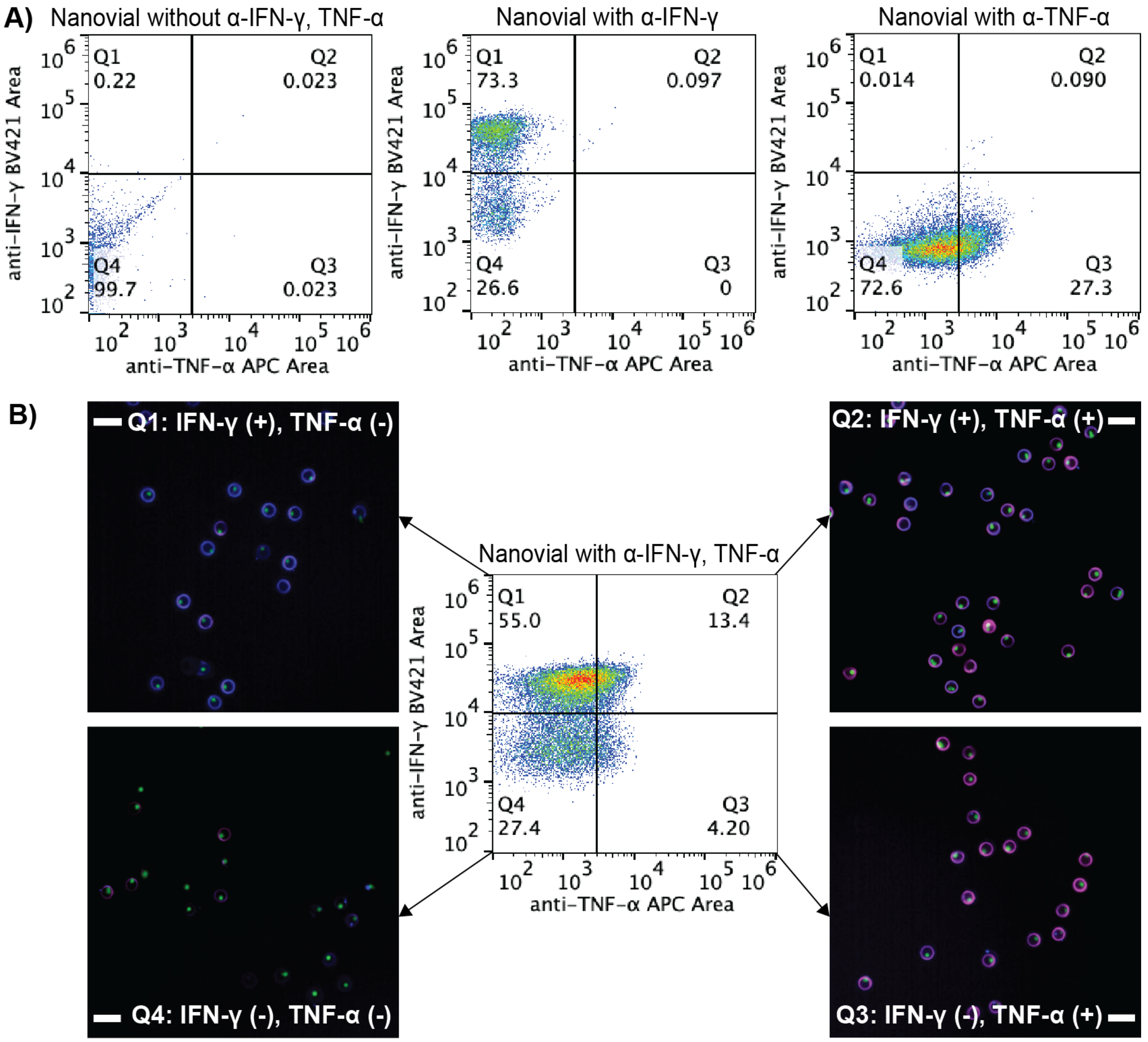


Figure S3 Analysis and sorting of single cells based on multiple secreted cytokines. A) Flow cytometry scatter plots of nanovials without any cytokine capture antibody and nanovials with only IFN-γ or TNF-α cytokine capture antibody are used to identify a positive threshold gate for IFN-γ, TNF-α or IFN-γ and TNF-α secreting cells. Each quadrant represents the following secreting population: Q1) IFN-γ secretors, Q2) IFN-γ and TNF-α polyfunctional secretors, Q3) TNF-α secretors, Q4) non-secretors. B) Flow cytometry scatter plots of T cells loaded onto nanovials labeled with all three antibodies (anti-CD45, anti-IFN-γ, anti-TNF-α) and sorted populations based on calcein signal and presence in one of the four quadrant gates according to each secretion phenotype. Green signal on microscopy images represents calcein signal. The majority of T cells heavily secreted IFN-γ and limited number of cells secreted TNF-α. Polyfunctional T cells (IFN-γ and TNF-α secretors) represented 13.4% of the population and were successfully sorted and recovered. Scale bars represent 50 µm.
